# NPM1 mediates genome-nucleolus interactions and the establishment of their repressive chromatin states

**DOI:** 10.1101/2024.07.03.601885

**Authors:** Shivani Gupta, Cristiana Bersaglieri, Dominik Bär, Mathieu Raingeval, Luana Schaab, Raffaella Santoro

## Abstract

Repressive chromatin domains are often located at the nuclear lamina (NL) or nucleolus. Although nucleolar associated domains (NADs) have been recently mapped, the mechanisms of NAD association with nucleoli and the functional significance of their localization remain unclear. Here, we show that NAD association with nucleoli is mediated by nucleophosmin (NPM1), a factor located within the granular component, the outer layer of the nucleolus. NPM1 binds NADs, interacts with the histone lysine methyltransferase G9a (EHMT2), and is required for establishing H3K9me2 at NADs. Loss of NPM1 or expression of NPM1 mutant lacking the DNA binding domain (NPM1_ΔDBD_) caused NAD dissociation from nucleoli and H3K9me2 reduction specifically at NADs. G9a is dispensable for NAD contacts with nucleoli and interacts with NPM1_ΔDBD_, indicating that NADs acquire G9a-mediated H3K9me2 only after associating with NPM1 at nucleoli. The results provide mechanistic insights into how genomic domains associate with nucleoli and acquire their repressive chromatin state. Additionally, our findings suggest that the nucleolus not only serves as a scaffold for positioning repressive chromatin domains but also plays a direct role in establishing these chromatin states.

## Introduction

In the nucleus of eukaryotic cells, chromosomes are arranged in a complex three-dimensional (3D) architecture that plays an important role to ensure the correct execution of gene expression programs. A key aspect of this regulation is the spatial positioning of genomic domains within the nuclear space. Generally, repressive chromatin regions are positioned at the nuclear lamina (NL) or nucleolus. Genomic regions interacting with NL are defined as lamina associated domains (LADs) whereas domains contacting nucleoli are named nucleolar associated domains (NADs) ^1,2^. LADs have been identified and characterized in many cell types and shown to be enriched in the histone marks H3K9me2/3 and H3K27me3 and harbour mostly inactive genes ^3,4^. The association of constitutive heterochromatin with the nucleolus has been documented by early microscopy studies ^5,6^. For example, in mouse cells, chromocenters, densely packed heterochromatic structures composed of highly repetitive pericentromeric sequences, are generally located close to nucleoli ^7^. However, only recently, the other non-repetitive fraction of NADs could be identified genome-wide in mammalian cells either by sequencing of biochemically purified nucleoli ^8–10^ or by applying DamID technology (Nucleolar-DamID)^11^. Although these techniques are based on different methodologies, they both revealed that NAD composition extends beyond centromeric or repetitive sequences and that NADs are generally in a repressive chromatin state ^12,13^. Moreover, the analysis of NADs in mouse embryonic stem cells (ESCs) identified three classes of genomic domains based on the interaction with nucleoli or NL: NAD-only that represent domains contacting the nucleolus but not the NL, LAD-only that contact the NL but not nucleoli, and NAD/LAD that can contact either nucleoli or the NL ^11,14^. The presence of NAD/LAD is also consistent with previous microscopy studies showing that some LADs can relocalize close to nucleoli after mitosis ^15^. Despite these advancements on the characterization of NADs, the mechanisms by which NADs associate with nucleoli and the functional significance of this interaction remain unclear. The nucleolus is a multilayer organelle that consists of the fibrillar center (FC) and the dense fibrillar component (DFC), surrounded by the granular component (GC). These nucleolar sub-compartments represent distinct, coexisting liquid phases ^16–18^. These structures play a major role in ribosome biogenesis, a process initiated by the RNA polymerase I (Pol I)-driven transcription of hundreds of ribosomal RNA (rRNA) genes that generates 45S/47S pre-rRNA ^19^. rRNA genes (rDNA) are organized in clusters and are distributed among different chromosomes. In humans and apes, they are located between the short arm and the satellite body of acrocentric chromosomes 13, 14, 15, 21, and 22. In mouse cells, rDNA repeats generally cluster close to the centromeric regions of chromosomes 12, 16, 18, and 19 (reviewed in ^2^). The linear genomic proximity of centromeric sequences to the rDNA locus explains why centromeres of chromosomes containing rDNA (rDNA-chromosomes) often cluster near nucleoli ^5^. However, centromeres from chromosomes not containing rDNA have also been shown to frequently localize near nucleoli ^6,11^, indicating that the contacts of centromeric regions with nucleoli is not only driven by their linear proximity to rDNA sequences. Several factors have been implicated in the proximity of centromeres or repetitive sequences to nucleoli, such as p150, a subunit of the Chromatin assembly factor-1 (CAF-1) in human cells ^20^ or Nucleolin homolog Modulo in *Drosophila* cells ^21^. However, it remains still elusive how the rest of the non-repetitive NADs are anchored to nucleoli and the functionality of these interactions.

The recent genome-wide identification of NADs in ESCs allowed us to finally investigate how these sequences associate with the nucleolus and the role of this interaction in terms of gene expression and chromatin states. Here we show that NAD association with nucleoli is mediated by components of the GC, NPM1 and Nucleolin, whereas the DFC component Fibrillarin is not involved in this process. We determined a direct role of NPM1 in NAD association with nucleoli and the establishment of histone lysine methyltransferase G9a (EHMT2)-mediated H3K9me2 at NADs. H3K9me2 represents the most prominent chromatin modification of NADs, forming a ring-like shape around nucleoli that could not be detected with any other repressive histone modifications. NPM1 associates with NADs and interacts with G9a. Loss of NPM1 or deletion of NPM1 DNA binding domain (DBD) caused NAD dissociation from nucleoli, the reduction of H3K9me2 specifically at NADs, and the disruption of the perinucleolar H3K9me2-marked ring. Remarkably, loss of NPM1 did not alter either the association of chromocenters with nucleoli or their characteristic H3K9me3 mark, indicating that NPM1 in ESCs only regulates the class of non-repetitive NADs. Finally, the data showed that G9a is dispensable for NAD contacts with nucleoli and interacts with NPM1_ΔDBD_, indicating that NADs acquire G9a-mediated H3K9me2 only after associating with NPM1 at nucleoli. The results provided mechanistic insights into how genomic domains associate with nucleoli and the establishment of their repressive chromatin structures. Moreover, they suggest that the nucleolus not only acts as a scaffold for the location of repressive chromatin domains but is also directly involved in the establishment of their chromatin states.

## Results

### GC components NPM1 and Nucleolin are required for NAD association with nucleoli

To determine which nucleolar components are implicated in NAD anchoring to nucleolus, we performed DNA-FISH in ESCs depleted of GC components NPM1 or Nucleolin or DFC component Fibrillarin via siRNA (**Fig. 1a-e**, **Extended Data Fig. 1**). We analysed four sequences corresponding to NAD-only regions at chromosomes 1, 2, 5, and rDNA-chromosome 19 that were previously identified and validated by Nucleolar-DamID ^11^ (**Fig. 1a**). These are strong NADs since they show contacts with nucleoli in more than 60-90% of ESCs (**Fig. 1e**). We found that the absence of GC components, in particularly NPM1, causes the detachment from nucleoli of the NADs of chromosomes 1, 2, and 5 (**Fig. 1b,e**, **Extended Data Figs. 1a-c**). The NAD of the rDNA-chromosome 19, which is close to the rDNA locus, remained attached to nucleoli in ESCs depleted of NPM1 or Nucleolin, an expected result since rDNA repeats should still be positioned within nucleoli (**Fig. 1e**, **Extended Data Figs. 1c,e**). In contrast, depletion of the internal DFC component Fibrillarin did not affect the location to nucleoli of all analysed NADs (**Fig. 1d,e**, **Extended Data Fig. 1f**). These results suggest that the GC compartment, which is the outer layer of the nucleolus, is implicated in NAD association with nucleoli.

**Figure 1.**
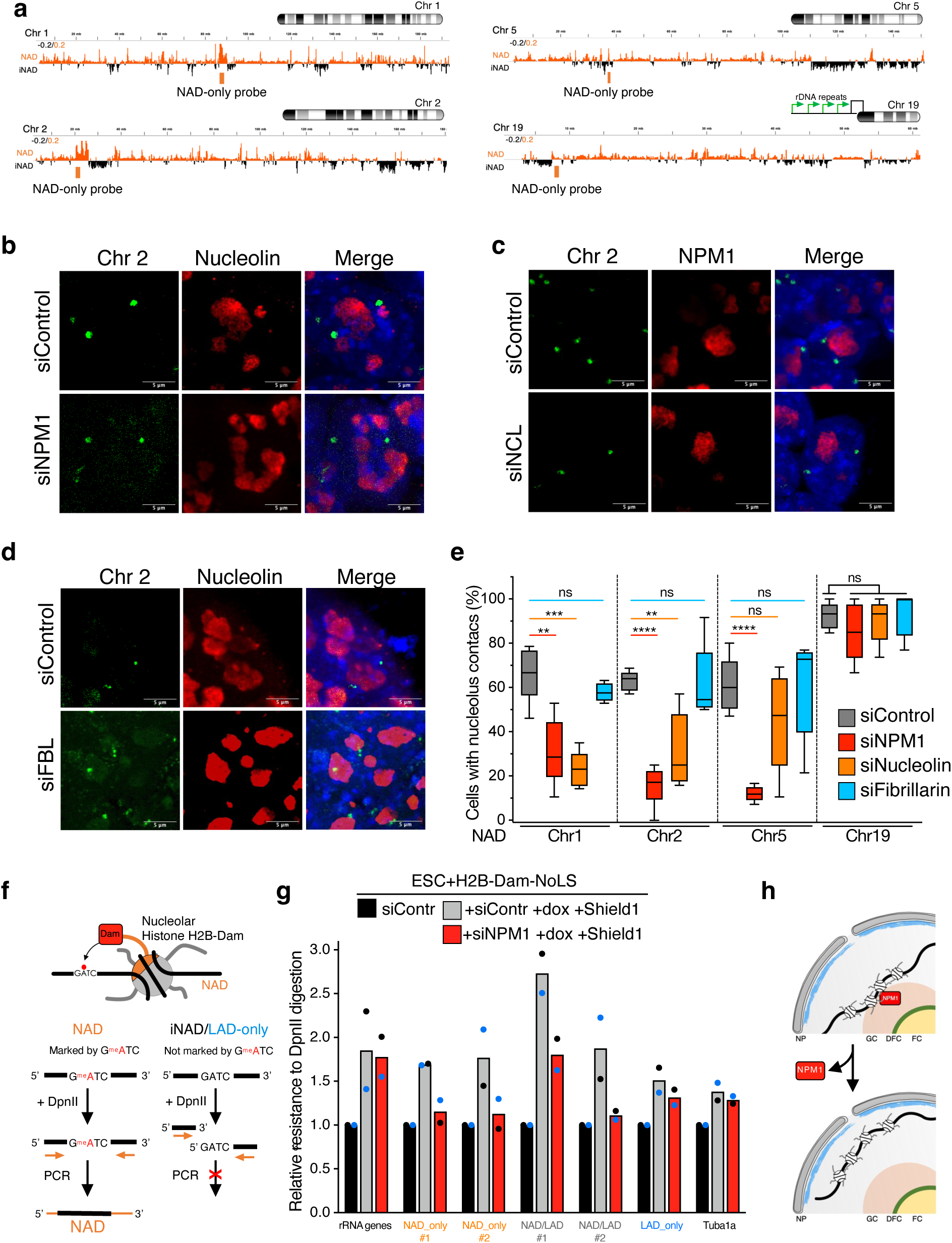
NPM1 and Nucleolin are required for NAD association with nucleoli. **a.** Representation of the DNA-FISH probes targeting NADs at chromosomes 1, 2, 5, and 19, the latter containing rRNA genes at the 5’, close to the centromeric region. The corresponding NADs profiles from Nucleolar-DamID ^11^ are shown. iNAD: genomic domains not interacting with nucleoli. **b-d.** Representative immuno-FISH images of NADs at chromosome 2 in ESCs depleted via siRNA of NPM1 (siNPM1, **b**), Nucleolin (siNCL, **c**), and Fibrillarin (siFBL, **d**). NPM1 and Nucleolin serve as nucleolar markers. **e.** Quantifications of immuno-FISH analyses showing the percentage of cells displaying NADs contacting nucleoli in ESCs depleted of NPM1, Nucleolin, or Fibrillarin. Error bars represent s.d. Statistical significance (*P*-values) from three independent experiments was calculated using the Mann-Whitney test (** < 0.01; *** < 0.001; ns: non-significant). **f.** Scheme representing the strategy to measure genomic contacts with nucleoli by Nucleolar-DamID coupled to DpnII digestion and qPCR. **g.** Relative resistance to DpnII digestion in ESCs expressing H2B-Dam-NoLS-DD with siRNA-Control siRNA-*NPM1*. Expression of H2B-Dam-NoLS-DD was induced by treatment with doxycycline and Shield for 15 hours. Data are from two independent experiments and values obtained in each experiment are shown as black or blue circles, respectively. The mean values are shown. **h.** Scheme representing the detachment of NADs from nucleoli upon loss of NPM1.

We focused our attention on NPM1 since the DNA-FISH analysis in NPM1-KD cells showed the strongest effect in the release of NADs from nucleoli. We validated these findings in an ESC line expressing NPM1 endogenously tagged with FKBP12^F36V^ that undergoes efficient protein degradation upon addition of dTAG-13 ^22^ (**Extended Data Figs. 2a-d**). The data also showed that loss of NPM1 does not affect the cell cycle (**Extended Data Figs. 2e**). We further supported the role of NPM1 in NAD association with nucleoli by performing Nucleolar-DamID coupled to qPCR in ESCs depleted of NPM1 (**Fig. 1f,g**). Nucleolar-DamID is a method that marks NADs via adenine methylation (^m6^A) at GATC sequences through the expression of an engineered nucleolar histone H2B fused to DNA adenine methyltransferase (H2B-Dam-NoLS) ^11^. NPM1-KD did not affect the nucleolar localization of H2B-Dam-NoLS, allowing the application of Nucleolar-DamID for NAD detection in NPM1 depleted cells (**Extended Data Fig. 2f**). ^m6^A levels were determined using the restriction endonuclease DpnII, which specifically cuts unmethylated GATC sequences, followed by quantitative amplification of NAD-only, NAD/LAD, and LAD-only sequences previously identified by Nucleolar-DamID ^11^. We used primers encompassing GATC sites to measure the amount of DpnII undigested DNA, which corresponds to sequences methylated by H2B-Dam-NoLS. Upon NPM1-KD, NAD-only and NAD/LAD sequences decreased m6A levels relative to control cells whereas m6A amounts at rRNA genes remained unaffected since they are still in contact with nucleoli (**Fig. 1g**). Moreover, LAD-only sequences and *Tuba1a*, which do not contact nucleoli, showed lower m6A levels relative to NADs and were not affected upon loss of NPM1, supporting the specificity of the Nucleolar-DamID. These results implicated an important role of NPM1 in mediating the association of NADs with nucleoli (**Fig. 1h**).

Chromocenters are often located close to nucleoli ^23^. These constitutive heterochromatic structures are composed of major and minor satellites and are highly enriched in the repressive chromatin mark H3K9me3 and heterochromatin protein 1 (HP1). Since a previous study in human cells reported that NPM1 associates with HP1 ^24^, we asked whether NPM1-mediated NAD anchoring to nucleoli could depend on HP1. We performed immuno-FISH on an ESC line engineered for 4-hydroxytamoxifen (4-OHT) inducible KO for HP1α and HP1β (*HP1α/β*-cDKO) ^25^ (**Extended Data Figs. 3a-c**). However, we did not find any evident alteration upon HP1α- and HP1β-KO, suggesting that NPM1-HP1 interaction is not required for anchoring NADs to nucleoli. Accordingly, the analysis of published ChIPseq data of HP1α, HP1β, and HP1γ in ESCs ^26^ revealed that the non-repetitive NADs are significantly depleted of HP1 relative to genomic domains located within the active A compartment (**Extended Data Fig. 3d**). Thus, while heterochromatic centric and pericentromeric satellite repeats are largely associated with HP1, the rest of the non-repetitive NADs are not significantly bound with HP1. Moreover, we did not observe any evident alteration in DAPI-stained heterochromatin organization around nucleoli upon HP1 loss, suggesting that HP1 is also not required for the association of chromocenters to nucleoli.

### NPM1 associates with NADs and regulates their G9a-mediated H3K9me2 content

Next, we asked whether NPM1 associates with NADs by performing NPM1-ChIPseq (**Fig. 2a**). The large majority of NPM1-bound chromatin corresponded to distal intergenic regions (65%) (**Fig. 2b**). We found that about one third of NPM1-bound regions correspond to NADs previously detected by Nucleolar-DamID whereas only a minor fraction (ca. 4%) are LAD-only sequences (**Fig. 2c**), indicating that NPM1 associates with genomic domains contacting the nucleolus but not with regions located at the nuclear periphery. To determine whether NPM1 regulates gene expression specifically at NADs, we performed RNAseq analysis of ESCs treated with siRNA-Control or siRNA-NPM1. We identified 239 upregulated genes and 471 downregulated genes in ESC depleted of NPM1 compared to control cells (log_2_ fold change ±0.58, *P* < 0.05, **Fig. 2d**). Genes significantly upregulated were enriched in pathways linked to cell differentiation and development whereas downregulated genes were implicated in diverse pathways (**Fig. 2e**). However, only a few genes at NADs (NAD-genes) were regulated by NPM1 (**Fig. 2f**), indicating that the detachment of genes from nucleoli upon NPM1-KD is not sufficient for their transcriptional activation.

**Figure 2.**
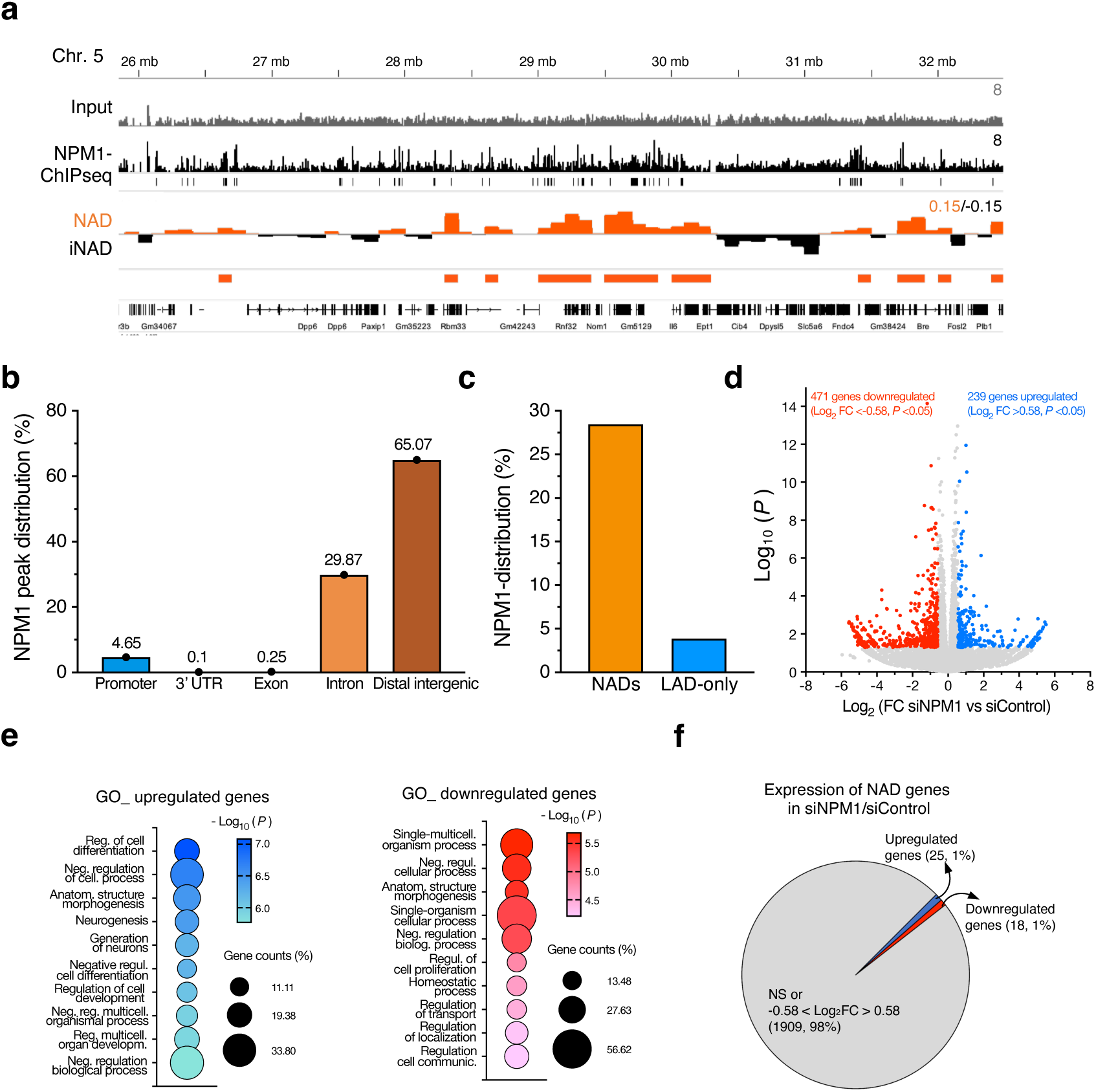
NPM1 associates with NADs. **a.** IGV track displaying NPM1 genome occupancy and NAD profiles from Nucleolar-DamID ^11^ of chromosome 5 in ESCs. **b.** Genomic annotations of NPM1-bound regions in ESCs. Promoter corresponds to ± 3kb regions from transcription start sites. **c.** NPM1 distribution on NADs and LAD-only sequences in ESCs. **d.** Volcano plot showing fold change (log_2_ values) in transcript levels of ESCs+siRNA-*NPM1* vs. ESC+siRNA-Control. Gene expression values of three replicates were averaged and selected for 1.5-fold changes and *P* < 0.05. **e.** Top 10 gene ontology (GO) terms as determined using DAVID for genes upregulated and downregulated upon NPM1-KD. **f.** Pie chart showing the number of genes located at NADs that are significantly up- or downregulated in ESC upon NPM1-KD.

Next, we analysed whether NPM1 could be implicated in the formation of repressive chromatin structure of NADs. Previous genomic studies in ESCs showed that NADs are enriched in H3K9me2 relative to genomic domains corresponding to the active A compartment whereas H3K9me3 levels were lower relative to A compartments and LADs ^11^. Consistent with these results, immunofluorescence analysis of ESCs showed the presence of H3K9me2-marked chromatin around nucleoli, forming a ring-like shape (**Fig. 3a,b**). These structures differed from H3K9me3-marked chromatin that corresponds to DAPI-stained heterochromatin and resembles chromocenters (**Extended Data Fig. 3e**). We determined that H3K9me2-ring around nucleoli is mediated by the euchromatic histone lysine methyltransferase 2 (G9a or EHMT2). Downregulation of G9a via siRNA significantly decreased H3K9me2 signal around nucleoli whereas depletion of three other H3K9 methyltransferases, Suvar39h1/2 and SETDB1, had no effect (**Fig. 3b,c**). Since this H3K9me2-marked chromatin around nucleoli could represent NADs, we asked whether NPM1, Nucleolin, or Fibrillarin regulate these structures. We performed H3K9me2 immunofluorescence and found that NPM1- or Nucleolin-KD cause strong alterations in the organization of H3K9me2-marked chromatin, as indicated by the loss of the H3K9me2 ring-like shape around nucleoli, whereas Fibrillarin-KD did not show any significant changes (**Fig. 3a,d**). Remarkably, NPM1 depletion did not cause any evident alterations in the nucleolar association of H3K9me3-marked and DAPI-stained heterochromatin and the corresponding major satellites sequences (**Extended Data Figs. 3e-g**). These results suggest that NPM1 regulates neither the nucleolar association of centromeric sequences nor their H3K9me3 content. Instead, NPM1 is required for the nucleolar anchoring of the large fraction of non-repetitive NAD sequences and the organization of the perinucleolar H3K9me2-marked chromatin.

**Figure 3.**
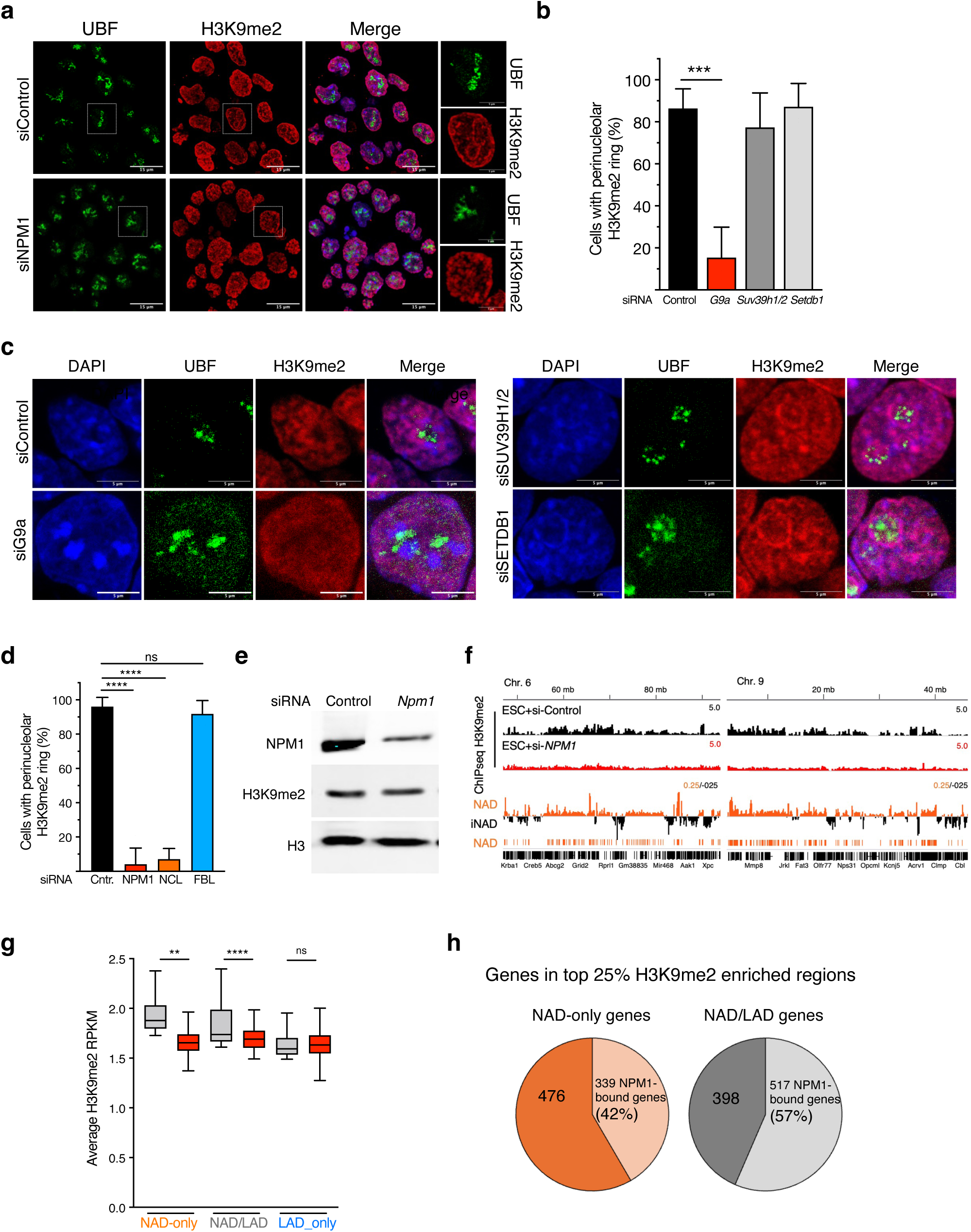
NPM1 regulates G9a-mediated H3K9me2 at NADs. **a.** Representative immunofluorescence images showing H3K9me2 distribution in ESCs treated with siRNA-Control or siRNA-*NPM1*. The magnification on the right shows the distribution of H3K9me2 around nucleoli, forming a ring-like shape, that is destroyed in the absence of NPM1. UBF serves as nucleolar marker. The corresponding DAPI staining is shown in **Extended Data Fig. 3f**. **b.** Quantification of cells showing the perinucleolar H3K9me2 ring in ESCs upon depletion of G9a, SUV39H1/2, or SETDB1. Data are from three independent experiments. Error bars represent s.d. Statistical significance (*P*-values) was calculated using the Mann-Whitney test (*** < 0.001). **c.** Representative immunofluorescence images showing H3K9me2 distribution in ESCs upon depletion of G9a, SUV39H1/2, or SETDB1 via siRNA. UBF serves as nucleolar marker. **d.** Quantification of cells showing the perinucleolar H3K9me2 ring in ESCs upon depletion of NPM1, Nucleolin (NCL), or Fibrillarin (FBL). Data are from three independent experiments. Error bars represent s.d. Statistical significance (*P*-values) was calculated using Mann-Whitney test (**** < 0.0001; ns: non-significant). **e.** Western blot showing global H3K9me2 levels in ESCs upon NPM1-KD. **f.** Tracks displaying H3K9me2 occupancy in ESCs treated with siRNA-Control or siRNA-NPM1 of chromosome 6 and 9 in ESCs. Lower panels show the corresponding NAD profile from Nucleolar-DamID ^11^. **g.** Quantifications of H3K9me2 levels at NAD-only, NAD/LAD, and LAD-only regions at 25% top H3K9me2 enriched regions in parental ESCs. Values from ESCs treated with siRNA-Control or siRNA-NPM1 are shown as average RPKM of a 10kb bin size region. Error bars represent s.d. Statistical significance (*P*-values) was calculated using the paired two-tailed t test (** < 0.01; ****<0.0001; ns: non-significant). **h.** Pie charts showing the number of genes located at the 25% top H3K9me2 enriched regions in parental ESCs and at NAD-only or NAD/LAD regions associated with NPM1.

Next, we asked whether the structural alterations of the perinucleolar H3K9me2-marked chromatin upon loss of NPM1 could reflect the detachment of NADs from nucleoli and/or alterations in H3K9me2 deposition. Western blot analysis revealed no evident changes in global H3K9me2 levels upon NPM1-KD (**Fig. 3e**). We performed quantitative H3K9me2-ChIPseq using spike-in chromatin in ESCs treated with siRNA-Control or siRNA-NPM1 (**Fig. 3f,g**). We found a significant downregulation of H3K9me2 at NAD-only and NAD/LAD regions that were enriched for this mark in parental ESCs (top 25%) whereas LAD-only did not show any significant changes, indicating a role of NPM1 in specifically regulating H3K9me2 at NADs. Moreover, we found that NPM1 associates with a large fraction of genes at NAD-only (42%) and NAD/LAD (57%) that are located at the top 25% enriched H3K9me2 chromatin domains (**Fig. 3h**). These results indicate that NPM1 directly regulates H3K9me2 content at NADs.

### NPM1 associates with G9a

To determine how NPM1 regulates H3K9me2 at NADs, we analysed whether NPM1 associates with G9a by performing NPM1-immunoprecipitation in ESCs followed by mass-spec analysis (**Fig. 4A**). The results showed that NPM1 associates with known nucleolar proteins, such as Fibrillarin and MKI67, and proteins highly enriched in functions related to ribosome biogenesis, nucleolus, gene expression, and chromatin organization (**Fig. 4b, Extended Data Fig. 4a**). Importantly, NPM1 also associates with both G9a and EHMT1 (GLP), which are known to form a stoichiometric heteromeric complex in a wide variety of cell types ^27^. We validated this interaction by HA-immunoprecipitation of HEK293T cells transfected with plasmids expressing HA-tagged G9a (**Fig. 4c**).

**Figure 4.**
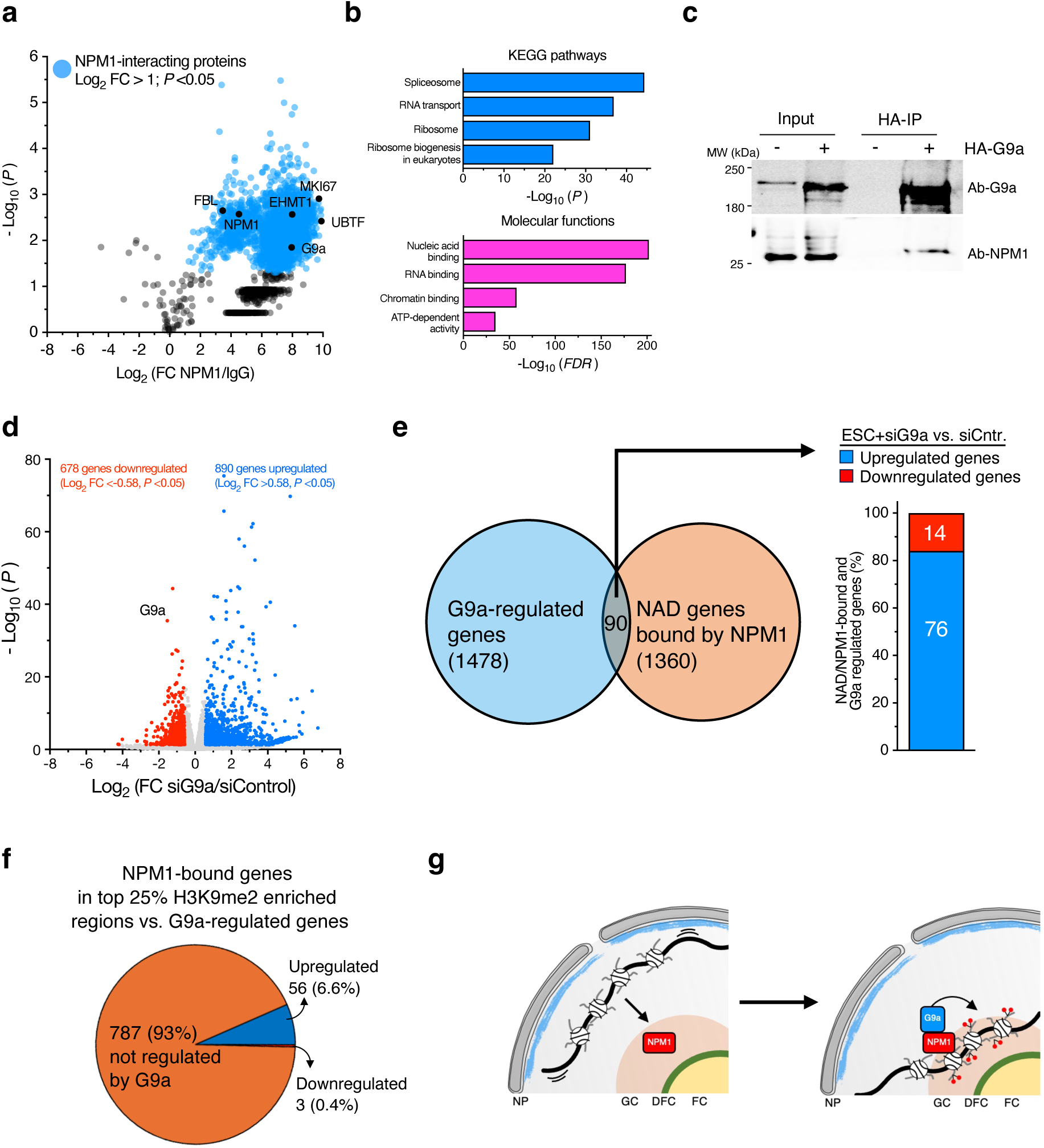
NPM1 associates with G9a. **a.** Volcano plot showing log_2_ fold changes of NPM1-immunoprecipitated peptides in ESCs relative to IgG control. Data are from three independent experiments. **b.** KEGG pathways and molecular functions of NPM1-interacting proteins detected by String analyses. **c.** HA-immunoprecipitation of HEK293T cells transfected with plasmid expressing HA-G9a. Signals were detected using antibodies against G9a and NPM1, respectively. **d.** Volcano plot showing fold change (log_2_ values) in transcript levels of ESCs+siRNA-*G9a* vs. ESC+siRNA-Control. Gene expression values of three replicates were averaged and selected for 1.5-fold changes and *P* < 0.05. **e.** Left panel. Venn diagram showing the number of G9a-regulated genes that are located at NADs and bound by NPM1. Right panel shows their changes in gene expression upon G9a-KD. **f.** Pie chart showing the numbers of NPM1-bound genes located at the top 25% of H3K9me2 enriched chromatin regions that are regulated by G9a. **g.** Scheme representing NMP1-mediated NAD association with nucleoli followed by NPM1-mediated recruitment of G9a to establish H3K9me2.

To determine whether H3K9me2 regulates the expression of genes located at NADs, we performed RNAseq of ESCs depleted of G9a via siRNA (**Fig. 4d**). As expected, G9a-KD strongly decreased global H3K9me2 levels (**Extended Data Figs. 4b,c**). Loss of G9a caused the upregulation of 890 genes and downregulation of 678 genes (log_2_ fold change ±0.58, *P* < 0.05, **Fig. 4d**). Genes significantly upregulated were enriched in pathways linked to cell adhesion whereas downregulated genes were mainly implicated in developmental processes (**Extended Data Fig. 4d**). However, we found that only few NAD-genes bound by NPM1 were regulated by G9a (90) and most of these genes (76%) were upregulated upon G9a-KD (**Fig. 4e**). Remarkably, the expression of NPM1-bound genes highly enriched in H3K9me2 was not affected by G9a-KD (93%, **Fig. 4f**). These results indicated that neither the detachment from nucleoli or the loss of G9a-mediated H3K9me2 is sufficient to derepress genes associating with nucleoli and/or NPM1 in ESCs. Next, we asked whether G9a-mediated H3K9me2 is required for NAD association with nucleoli. We performed DNA-FISH and Nucleolar-DamID analyses in ESCs depleted of G9a via siRNA (**Extended Data Figs. 4e-g**). Both analyses showed that NAD location at nucleoli was not altered in the absence of G9a, indicating that H3K9me2 does not regulate NAD association with nucleoli. Together, these results suggest that once NADs are anchored at nucleoli through NPM1 binding, NPM1 recruits G9a to establish H3K9me2 at these chromatin domains (**Fig. 4g**). Thus, the establishment of repressive chromatin at NADs can only occur when NADs interact with NPM1 and associate with nucleoli.

### NPM1-mediated NAD association with nucleoli depends on the DNA-binding domain of NPM1

To determine how NPM1 mediates the association of NADs with nucleoli, we analysed the impact of several known features of NPM1. NPM1 is known to form oligomers through its N-terminal domain ^28,29^. These NPM1 structures seem to contribute to the liquid-like features of the GC compartment through both heterotypic and homotypic LLPS mechanisms ^30^. To determine whether NPM1 oligomerization properties are important for NAD association with nucleoli, we treated ESCs with the small molecule NSC348884 that was reported to prevent formation of NPM oligomers ^31^. Accordingly, treatment for 24 hours with NSC348884 concentrations ranging from 0.3 to 1.0 μM strongly impaired the formation of NPM1 oligomers without affecting NPM1 nucleolar localization (**Extended Data Figs. 5a,b**). However, treatments longer than 24 hours were very toxic to cells, leading to their detachment from the plate. Thus, for downstream analyses, we used conditions that impair NPM1 oligomerization without severely affecting cell viability (i.e., 0.5 μM for 24 hours). Immuno-FISH and immunofluorescence analyses did not detect significant changes in NAD contacts with nucleoli or alterations of the H3K9me2 perinucleolar ring between treated and untreated cells (**Fig. 5a-c**, **Extended Data Fig. 5c**), indicating that NPM1 oligomerization might not contribute to NAD association with nucleoli and the establishment of H3K9me2 at NADs.

**Figure 5.**
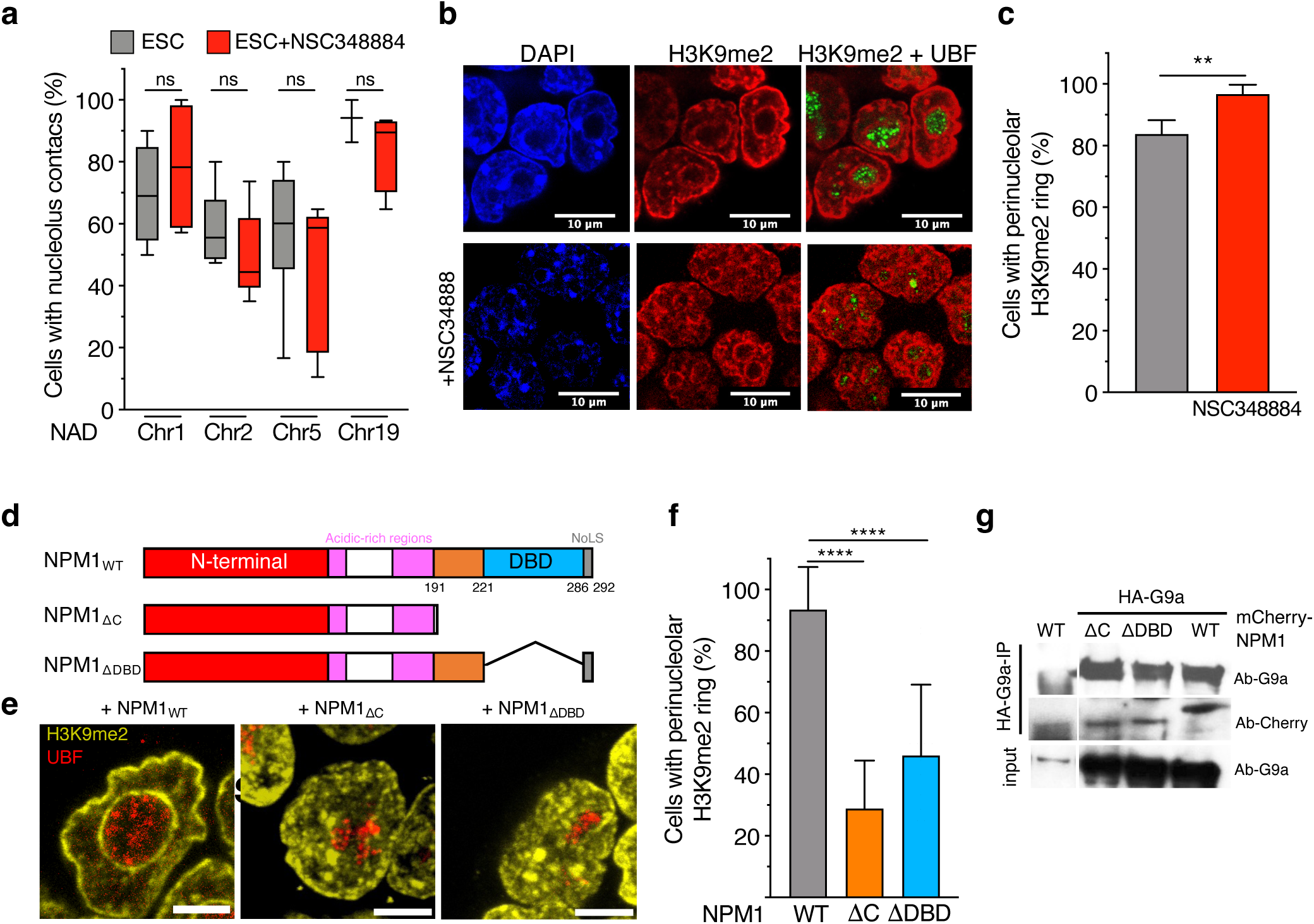
NPM1 C-terminal domain is required for H3K9me2 chromatin organization around nucleoli. **a.** Quantifications of immuno-FISH analyses showing the percentage of cells displaying NADs contacting nucleoli in ESCs treated with NSC348884. Error bars represent s.d. Statistical significance (*P*-values) from three independent experiments was calculated using the Mann-Whitney test (ns: non-significant). **b.** Representative H3K9me2 immunofluorescence images of in ESCs treated for 24 hours with 0.5 μM NSC348884. UBF serves as nucleolar marker. **c.** Quantification of cells showing the perinucleolar H3K9me2 ring in ESCs upon treatment with NSC348884. Error bars represent s.d. Statistical significance (*P*-values) from three independent experiments was calculated using the Mann-Whitney test (**: <0.01). **d.** Scheme representing the NPM1 mutants and the corresponding domains. **e.** Representative H3K9me2 immunofluorescence images of ESCs expressing NPM1_wt,_ NPM1_ΔC_, or NPM1_ΔDBD_. UBF serves as nucleolar marker. Scale bar is 5 μm. **f.** Quantification of the perinucleolar H3K9me2 ring in ESCs expressing NPM1_wt,_ NPM1_ΔC_, or NPM1_ΔDBD_. Error bars represent s.d. Statistical significance (*P*-values) from three independent experiments was calculated using the Mann-Whitney test (****: <0.0001). **g.** HA-immunoprecipitation of HEK293T cells co-transfected with plasmid expressing HA-G9a and mCherry-NPM1 constructs. Signals were detected using antibodies against G9a and mCherry, respectively.

Next, we generated NPM1 mutants by deleting the C-terminus part that contains both the DNA binding domain (DBD) and the nucleolar localization signal (NoLS) (NPM1_ΔC_) or only the DBD (NPM1_ΔDBD_) (**Fig. 5d**). We found that the expression of both NPM1 mutants in ESCs alters the H3K9me2-marked nucleolar ring whereas the expression of NPM1_WT_ did not show any alterations (**Fig. 5e,f**). Given the presence of endogenous NPM1, these results also indicated that NPM1 mutants act in a dominant negative manner, as also reported for the C-terminus mutated NPM1 (NPM1c) in acute myeloid leukemia (AML) ^32^. Notably, immunoprecipitation analyses in HEK293T cells showed that both NPM1 mutants can still associate with G9a (**Fig. 5g**), suggesting that G9a interacts with the N-terminal NPM1 domains and that the establishment of H3K9me2 at NADs requires NPM1 association with NADs via its DBD. Together, the results showed that NADs acquired G9a-mediated H3K9me2 only when associating with NPM1 at nucleoli, indicating that the nucleolus does not only act as a scaffold for the location of repressive chromatin domains but is also directly involved in the establishment of these chromatin states.

## Discussion

Microscopy and recent genomic data have highlighted the nucleolus as a compartment where repressive chromatin domains are often located. However, how NADs associate with nucleoli and whether and how this interaction is involved in the establishment or maintenance of repressive chromatin and gene expression states has remained elusive.

Here we have analysed mouse ESCs taking the advantage that the non-repetitive fraction of NADs in ESCs has been recently mapped genome-wide ^11^. We show that the outer layer of the nucleolus, the GC compartment, is required for NAD association with nucleoli. In particular, we show that NPM1 binds to NADs and stabilizes their association with nucleoli. Importantly, we showed that NPM1 does not regulate the nucleolar association of constitutive heterochromatic sequences such as centromeric repeats, which are characterized by H3K9me3 and positive to DAPI staining. This result could be in part explained since centromeric sequences from rDNA-chromosomes are in linear proximity to rDNA repeats and consequently remained localized at nucleoli upon NPM1 loss. Moreover, our data indicated that NPM1 does not affect the organization of H3K9me3-marked chromatin that is highly enriched at centromeric sequences ^33^. An early microscopy study in human cells has shown that NPM1-KD alters the distribution of DAPI-stained and H3K27me3 and H3K9me3 heterochromatin around nucleoli ^24^. Although this study did not directly address whether these changes correspond to a loss of NAD association with nucleoli, it suggested together with our work that the features of heterochromatin around nucleoli might differ between mouse and human cells. Accordingly, we showed that the nucleoli of mouse ESCs display a ring of chromatin enriched in H3K9me2 and negative to DAPI-staining whereas H3K9me3 forms large bodies corresponding to chromocenters that are adjacent to nucleoli without surrounding them. In contrast, human cells do not show prominent chromocenters ^34^ and nucleoli are generally encircled by DAPI-stained heterochromatin. Thus, microscopy data suggest that a large fraction of NADs in human cells are more enriched in constitutive heterochromatin than in mouse cells, explaining why loss of NPM1 has different effects on constitutive heterochromatin at nucleoli between mouse and human cells.

The analyses of NADs in ESCs indicated that NPM1 regulates the establishment of H3K9me2 at NADs. Our results showed that the nucleolus is surrounded by H3K9me2-marked chromatin, while other repressive histone modifications were not detected. This is supported by previous work demonstrating that H3K9me2 is the predominant histone modification at non-repetitive NADs, with higher levels than in the active A compartment and lower H3K9me3 and H3K27me3 levels compared to LAD-only regions and active chromatin ^11^. Mechanistically, we showed that NPM1 recruits G9a at NADs to establish H3K9me2. Moreover, we showed that G9a-mediated H3K9me2 at NADs depends on NPM1, suggesting that this modification occurs at nucleoli. Accordingly, the expression NPM1 mutant lacking DBD destroys the H3K9me2 ring but its interaction with G9a was not affected. These results suggested that the nucleolus acts as regulator for the establishment of repressive chromatin domains, extending beyond its known role to serve as scaffold for positioning repressive chromatin domains in the nuclear space. Nevertheless, at least in ESCs, neither the KD of NPM1 nor G9a reactivates gene expression at NADs, suggesting that the detachment from nucleoli or the loss of repressive H3K9me2 mark is not sufficient for gene reactivation. However, ESCs might represent a special case since it is well known that loss of repressors, such as DNA methyltransferases, have little effect on gene expression ^35^. Our results also might have an impact on human diseases linked to nucleolar alterations such as AML where the nucleolar function of NPM1 is impaired in 35% of AMLs as a result of *NPM1c* mutations ^36^, which disrupt the N-terminal nucleolar localization signal and generate a nuclear export signal in its place ^37^. Understanding the impact of potential alterations in genome organization and chromatin state around nucleoli in AML cells harboring NPM1c paves the way for future studies.

Together, our findings reveal the mechanisms by which genomic domains associate with nucleoli and how their repressive chromatin structures are formed. Moreover, they indicate that the nucleolus not only acts as a scaffold for positioning repressive chromatin domains but is also directly involved in the formation of their repressive state.

## Materials and methods

### Cell culture

One hundred and twenty-nine mouse embryonic stem cells (E14 line) were cultured in 2i media composed of DMEM-F12 (ThermoFisher Scientific, 11320) and Neurobasal medium (ThermoFisher Scientific, 21103) in 1:1 ratio, supplemented with 1× N2/B27 (Gibco 17502, 17504), 1× penicillin/streptomycin/l-glutamine (Gibco, 10378), 50 μM β-mercaptoethanol (Gibco, 31350), recombinant leukemia inhibitory factor, LIF (Polygene, 1,000 U/ml) and MEK and GSK3β inhibitors, 2i (Sigma CHIR99021 and PD0325901, 3 and 1 μM, respectively). ESCs were seeded at a density of 50,000 cells/cm^2^ in culture dishes (Corning^®^ CellBIND^®^ surface) coated with 0.1% gelatin (Sigma, G1890) without a feeder layer. Propagation of cells was carried out every 2 days using Trypsin-EDTA (Gibco, 15400) mediated cell dissociation. *HP1α/β*-cDKO ESC line is a kind gift from Antoine H.F.M Peters lab ^25^. They were grown in same manner as E14 cell line.

HEK 293T and NIH3T3 cells were cultured in Dulbecco’s modified Eagle’s medium (ThermoFisher Scientific, 61965) supplemented with 10% fetal calf serum (FCS, Biowaste) and 1% penicillin/streptomycin (Gibco, 15140).

### Transfections

ESCs were transfected with the indicated siRNAs (40 nM siRNA if not stated differently) using Lipofectamine® RNAiMAX Transfection Reagent (Invitrogen, 13778150) in Opti-MEM® GlutaMAX™ (Gibco, 31985062) reduced-serum medium Transfection Reagent (Invitrogen 13778150) following the manufacturer’s instructions. 8×10^4^ parental ESCs were seeded into gelatin-coated 6-well plate (Corning^®^ CellBIND^®^ surface) 4 hours prior to siRNA transfection and collected 3 days after transfection. In the case of Nucleolin, we performed two consecutive treatments with siRNA-control and siRNA-Nucleolin (80nM siRNA), each one lasting for 3 days. Efficiencies of siRNA-mediated depletions were monitored by qRT-PCR and western blot. Only samples showing reduction levels up to 70-80% were processed for further downstream analyses. For DNA FISH, 20×10^3^ ES cells were plated per well of Matrigel (Corning, 356238) coated Nunc Lab-Tek II chamber Slide System (Thermo Fisher Scientific, 154526) 4 hours prior to siRNA transfection as described above. For H3K9me2-ChIP, 0.4×10^6^ ES cells were seeded into gelatin coated 10 cm dishes 4 hours prior to siRNA transfection.

For transient transfection of NIH3T3 cells, 1.5×10^5^ cells were plated in 6-well plates and transfected respectively with 200ng of plasmid expressing H2B-GFP or H2B-GFP-NoLS under the control of minimal CMV promoter, and with 1.5μg of plasmid expressing shRNA-Control or shRNA-*Npm1* using Transit-X2 transfection reagent (Mirus) in Opti-MEM GlutaMAX reduced-serum medium (Life Technologies). 72h post-transfection, cells were fixed for immunofluorescence.

For transient transfection of HEK 293T cells, 2.5×10^6^ cells were plated on 15cm dishes. The day after cells were transfected with the indicated plasmids using BES-CaCl_2_ transfection method. After 48 hours cells were collected for immunoprecipitation experiments.

### Generation of FLAG-HA tagged and FKBP-tagged NPM1 cell line

CRISPR/Cas9 cloning and targeting strategy were performed as previously described. ESCs were co-transfected with a plasmid expressing Cas9 protein and the gRNA sequence targeting the NPM1 locus (pX330 Mouse 5’ Npm1 gRNA, a gift from Mark Groudine, Addgene plasmid #127900; http://n2t.net/addgene:127900; RRID:Addgene_127900) and the HDR donor plasmid derived from Mouse 5’ Npm1-AID-GFP-PuroR plasmid (a gift from Mark Groudine, Addgene plasmid #127899 ;http://n2t.net/addgene:127899; RRID:Addgene_127899) in which AID-GFP sequences were replaced with FLAG-HA or FKBP sequence (synthesized from IDT). The transfection was done with TransIT-X2 transfection reagent (Mirus Bio, MIR6004) following the manufacturer’s instructions. 15×10^4^ wild-type ESCs were seed into gelatin-coated 6 cm plate, let to attach for 4 hours, and then transfected. Two days after transfection, ESCs were selected using 2 μg of Puromycin (Gibco, A11138) overnight. After recovery, ESCs were further treated with 1 μg of Puromycin for additional three days. After three days of culture, cells were seeded for single cell clone isolation. Puromycin resistant ESC clones were analysed by western blot to identify cells expressing NPM1 in one or both alleles. ESC with homozygous insertion were selected for all the analyses.

### DNA-FISH

DNA FISH was performed as previously prescribed ^11^. Briefly, ESCs were grown on matrigel coated Nunc Lab-Tek II chamber Slide System (Thermo Fisher Scientific, 154526) for all the DNA FISH assays. For Actinomycin D treament, 20×10^3^ cells were grown per well of Lab-Tek II chamber slides for 24 hours prior to the treatment. *HP1α/β*-cDKO cells were treated with 1µM 4-Hydroxytamoxifen (Sigma, H6278) for 48 hours. After the treatment/transfections, cells were fixed with 4% PFA for 10 minutes at room temperature. Cells were washed three times with 1x PBS and then permeabilized with 0.5%Triton-×100/PBS (Triton-X 100: Sigma, T8787) for 10 min at room temperature. After three additional washes with 1X PBS, cells were incubated with 0.1 M HCl for 10 min at room temperature, washed twice with 2x SCCT, and twice with 50% Formamide-2X SCCT (Formamide: Sigma, 47671). Probe mixture contains Xpmol of Oligopaint probe, 0,8uL of ribonuclease A (ThermoFisher Scientific, EN0531, 10 mg/mL), and FISH hybridization buffer for a total mixture volume of 10uL. The probe mixture was added directly on the chamber slide. The cellular DNA was denatured at 83 °C for 8 min, and hybridization was performed in a humid dark chamber overnight at 42 °C. Cells were washed twice with 2X SCCT for 15 min, once with 0.2X SCC for 10 min, twice with 1X PBS for 2 min, and three times with 1X PBT for 2 min. Slides were then incubated for 1 h at room temperature with blocking solution in a dark humid chamber, and overnight at 4 °C with primary antibody against Nucleolin or NPM1 (Nucleolin - abcam, ab22758 1:1000; NPM1 – Sigma, B0556 1:500). Cells were washed three times with 1X PBT for 5 minutes each and then incubated for 1 h at room temperature with secondary antibody (goat anti-rabbit IgG (H+L) Alexa Flour 546 (Invitrogen, A11035) or goat anti-mouse IgG (H + L) Alexa Fluor 546 (Invitrogen, A11030)) in a dark humid chamber. The slides were washed three times with 1X PBT for 5 minutes each. Then, the slides were mounted with Vectashield DAPI mounting media (Vector, H-1200) and, after drying, stored at 4 °C.

Immuno-DNA-FISH samples were imaged using a Leica SP8 upright Microscope, with a z-stack collected for each channel (step size, 0.15 or 0.3 um, frame interval 1 s), using the oil objective HC PL APO CS2 63x/1.40. Images were processed by ImageJ (version 2.0.0/1.53c). The individual cells were identified by Hoechst/DAPI staining and cells containing signal for DNA-FISH channel were identified manually on the corresponding fluorescent channel. Distance between the DNA-FISH signal and the nucleolar marker immunofluorescence signal was calculated using ImageJ (version 2.0.0/1.53c) and used to count the number of foci contacting nucleolus and the number of cells with at least one contact with the nucleolus. DNA-FISH probes for chromosomes 1, 2, 4, 5, and 19 were generated with oligopaint libraries that were constructed the PaintSHOP interface created by the Beliveau lab (https://oligo.shinyapps.io/paintshop/_w_33571817/#tab-1201-8) and were ordered from CustomArray/Genscript in the 12 K Oligopool format. Each library contains a universal primer pair used to amplify all the probes in the library, followed by a specific primer pair hooked to the 40–46-mer genomic sequences, for a total probe of around 124-130-mers. Oligopaint libraries were produced by emulsion PCR from the pool, followed by a “two-step PCR” and lambda exonuclease as described before. Specifically, the emulsion PCR with the universal primers allowed amplifying all pool of probes in the library. The “two-step PCR” led to the addition of a tail to the specific probes bound to the Alexa Fluor 488 fluorochrome. All oligonucleotides were purchased from Integrated DNA Technologies (IDT, Leuven, Belgium). The sequence of DNA FISH probe for major satellite was taken from previously published work ^38^. The probe of major satellite was labelled at 5’ end Californian Red and synthesized from Microsynth.

### Immunofluorescence

Cells were plated on coverslips (VWR, ECN 631-1577) and pre-coated with Matrigel (Corning, 356238) for 1 hour at 37°C. After 48 to 72 hours, depending on the treatment, cells were fixed using 3.7% formaldehyde (Sigma-Aldrich, 47608) in PBS for 10 minutes at room temperature. After two washes in PBS, cells were permeabilized with 0.5% TritonX-100 (Sigma-Aldrich, T8787) in PBS for 10 minutes on ice and 10 minutes at room temperature. Cells were then washed two times in PBS and blocked with 1% Bovine Serum Albumin, (Sigma-Aldrich, A2153) in PBS-0.1% Tween 20 (Tween 20: Merck, P9416) for 1 hour at room temperature in a humidified chamber. The coverslips were then incubated with primary antibodies resuspended in blocking solution overnight at 4°C in a humidified chamber. After 3 washes in 1XPBS, the coverslips were incubated with secondary Alexa Fluor^®^ antibody resuspended 1:1000 in Blocking Solution for 1 hour at room temperature. After 3 washes in 1xPBS, the coverslips were mounted with VECTASHIELD^®^ Antifade Mounting Medium with DAPI (H-1200-10), sealed with nail polish and imaged within one week.

### Image acquisition and analysis

Images were acquired with a Leica inverse SP8 FALCON (Fast Lifetime CONtrast) confocal laser scanning microscope, equipped with 63x HCX PL APO CS2 objective with oil immersion. For quantification a Z-stack (10-15 x 0.3 μm step size) of 15 to 20 colonies were imaged for each condition. The images were processed using FIJI (version 2.14.0/1.54f).

### Nucleolar-DamID

5 x 10^4^ H2B-Dam-NoLS-DD ESCs ^11^ were seeded in 6-well plates (Corning^®^ CellBIND^®^ surface) coated with 0.1% gelatin without feeder layer. Cells were left to attach for 4h and then transfected with either siRNA targeting *Npm1*, *G9a*, or Control using Lipofectamine RNAiMAX transfection reagent (Invitrogen, 13778150) following the manufacturing instructions. 3 days after siRNA transfection, cells were treated with 100ng/mL Doxycycline and 1µM Shield1 (Clontech Takara) for 15h to induce the expression of Dam-fused proteins. Cells were harvested by trypsinization, and DNA was extracted using Quick-DNA Miniprep Plus kit (Zymo Research). m6A levels at GATC were measured by Dpn II digestion followed by quantitative PCR.

### Cell cycle analysis

The FKBP12F36V-NPM1-ESCs were treated with dTAG-13 (R&D Systems, 6605/5) at 100nM for 3 days and DMSO was used as a control. Thereafter, the cells were trypsinized and washed with cold 1X PBS. Cells were fixed with cold 70% ethanol and stained with propidium iodide at a final concentration of 20 μg/ml (Sigma, P4864) at room temperature for 30 minutes in dark. The cell cycle data was collected by flow cytometry FACS canto II (BD Biosciences). The cell cycle data was analysed by using the software FlowJo (v10.10).

### RNA extraction, reverse transcription, and quantitative PCR

RNA was purified with TRI reagent (Molecular Research Center, Inc., TR118). 1 µg total RNA was primed with random hexamers and reverse-transcribed into cDNA using MultiScribe™ Reverse Transcriptase (Invitrogen, 4311235). Amplification of samples without reverse transcriptase assured the absence of genomic or plasmid DNA. The relative transcription levels were determined by normalization to 𝛽*-Actin* mRNA levels. qRT-PCR was performed with KAPA SYBR^®^ FAST (Sigma) on a Rotor-Gene Q (Qiagen).

### RNAseq and data analysis

Total RNA was purified with TRI reagent (Molecular Research Center, Inc., TR118). The quality of the isolated RNA was determined with a Qubit^®^ 4 Fluorometer and 4200 Tape Station system. Only those samples with a 260 nm/280 nm ratio between 1.8–2.1 and a 28S/18S ratio within 1.5-2 were further processed. The TruSeq Stranded mRNA kit (Illumina) was used for library preparation in the following steps. Briefly, total RNA samples (100-1000 ng) were polyA enriched and then reverse-transcribed into double-stranded cDNA. The cDNA samples were fragmented, end-repaired and adenylated before ligation of TruSeq adapters containing unique dual indices (UDI) for multiplexing. Fragments containing TruSeq adapters on both ends were selectively enriched with PCR. The quality and quantity of the enriched libraries were validated using Qubit^®^ 4 Fluorometer. The product is a smear with an average fragment size of approximately 260 bp. Libraries were normalized to 10 nM in Tris-HCl 10 mM, pH8.5 with 0.1% Tween 20. The Illumina NovaSeq 6000 was used for cluster generation for RNAseq of NPM1 knockdown and sequencing performed was single end 100bp according to standard protocol. For RNAseq of ESCs upon G9a knockdown, Illumina NovaSeq X Plus was used for cluster generation and sequencing was paired end at 2×150bp according to standard protocol. The quality of the reads generated by the machine was checked by FastQC, a quality control tool for high throughput sequence data. The quality of the reads was increased by applying: a) SortMeRNA ^39^ (version 2.1) tool to filter out ribosomal RNA; b) Trimmomatic ^40^ (version 0.40) software package to trim the sorted (a) reads. The sorted (a), trimmed (b) reads were mapped against the mouse genome (mm10) using the default parameters of the STAR (Spliced Transcripts Alignment to a Reference, version 2.7.0a) ^41^. For each gene, exon coverage was calculated using a custom pipeline and then normalized in reads per kilobase per million (RPKM) ^42^, the method of quantifying gene expression from RNA sequencing data by normalizing for total read length and the number of sequencing reads.

### ChIPseq and data analysis

ChIP analysis was performed as previously described ^43^. For H3K9me2 ChIPseq, parental ESCs were used. For NPM1-ChIPseq, the ESC lines expressing endogenous NPM1 tagged with FLAG-HA was used. Briefly, 1% formaldehyde was added to cultured cells to cross-link proteins to DNA. Isolated nuclei were then lysed with lysis buffer (50 mM Tris-HCl, pH 8.1, 10 mM EDTA, pH 8, 1% SDS, 1X cOmplete™ protease inhibitor EDTA-free cocktail (Roche, 1187358001). Nuclei were sonicated using a Bioruptor ultrasonic cell disruptor (Diagenode) to shear genomic DNA to an average fragment size of 200 bp. 20 μg of chromatin for H3K9me2 ChIP and 50ug of chromatin for NPM1 CHIP was diluted to a total volume of 500 μl with ChIP buffer (16.7 mM Tris-HCl, pH 8.1, 167 mM NaCl, 1.2 mM EDTA, 0.01% SDS, 1.1% Triton X-100) and pre-cleared with 10 μl packed Protein A Sepharose CL-4B resin (Cytivia, 17078001) for 2 hours at 4°C. Pre-cleared chromatin was incubated overnight with the ChIP-grade antibodies against H3K9me2 (abcam, ab1220) and HA (abcam, ab9110). The next day, Dynabeads™ Protein A (Invitrogen, 10001D) for HA-ChIP or Protein G (Invitrogen, 10003D) for H3K9me2-ChIP were added and incubated for 4 hours at 4°C. After washing, bound chromatin was eluted with the elution buffer (1% SDS, 100 mM NaHCO_3_). Upon proteinase K (ThermoFisher Scientfic, EO0491) digestion at 50 °C for 3 h and reversion of cross-linking (65 °C, overnight), DNA was purified with phenol/chloroform, ethanol precipitated and quantified.

ChIP-qPCR measurements were performed with KAPA SYBR® FAST (Sigma) on a Rotor-Gene Q (Qiagen) always comparing enrichments over input samples.

For ChIPseq library preparation, the quantity and quality of the isolated DNA was determined with Qubit® 4 Fluorometer (Life Technologies). Libraries were prepared using the NEBNext^®^ Ultra^™^ II DNA Library Prep for Illumina (New England Biolabs, E7645S and E7645L) following the manufacturer’s protocol. Briefly, ChIP and input samples (10 ng) were first end-repaired and polyadenylated. Then, the ligation of Illumina compatible adapters containing the index for multiplexing was performed. The quality and quantity of the enriched libraries were evaluated using Qubit^®^ 4 Fluorometer and 4200 TapeStation System (Agilent). Sequencing was performed on an Illumina NovaSeq6000 machine with single-end 100 bp reads for NPM1 and H3K9me2 according to standard protocols.

The ChIPseq reads were aligned to the mouse mm10 reference genome using Bowtie2 (version 2.3.4.3). Read counts were computed and normalized using “bamCoverage” from deepTools (version 3.2.1) using a bin size of 50 bp. To calculate read coverage for 20 kb bin region of H3K9me2 ChIPseq, “multiBamSummary” from deepTools was used. NPM1 bound regions were defined using MACS2 by comparing the HA CHP of tagged NPM1 ESCs and input. HP1α and HP1ý CHIPseq data were from a previous report ^26^ and publically available on GEO, GSE97945. To calculate read coverage for 50 kb bin region of ChIPseq HP1α and HP1ý, “multiBamSummary” from deepTools was used. Integrative Genome Viewer (IGV, version 2.5.2) was used to visualize and extract representative ChIPseq tracks.

### NSC348884 treatment

48 hours post seeding, ESCs were treated with NSC348884 (Selleckchem,S8149) for 24 hours NSC348884. Cell viability was monitored by bright field microscopy. For immunofluorescence and DNA-FISH, cells were fixed with 3.7% formaldehyde. For protein analysis, cells were trypsinized and centrifuged at 210 g for 5 min. 1×10^6^ cells were resuspended in 50 μl lysis buffer (10 mM Tris-HCl pH 7.5, 150 mM NaCl, 0.5 mM EDTA, 0.5% NP-40) and 2X native buffer (50 mM Tris-HCl, 10 mM dithiothreitol (Sigma, D9779), 10% glycerol (Sigma, 49781) was added to the lysate in a 1:1 ratio.

### Immunoprecipitation

For NPM1-IP, cells were collected and the nuclei were isolated as previously described ^44^. Immunoprecipitation analysis was performed with 10 x 10^6^ parental ESCs. Cells were washed twice in cold 1X PBS and then incubated with hypotonic buffer (10mM HEPES, 1.5mM MgCl_2_, 10nM KCl, 1X protease inhibitor) for 10 minutes on ice. Samples were supplemented with Triton X-100 at a final concentration of 0.5% for 10 minutes on ice. Then, cell nuclei were lysed for 30 minutes at 37°C in MNase digestion buffer (50 mM Tris–HCl pH 7.5, 0.3 M Sucrose, 30 mM KCl, 7.5 mM NaCl, 4 mM MgCl_2_, 1 mM CaCl_2_, 0.125% NP-40) freshly supplemented with 0.25% NaDeoxycholate, cOmplete™ Protease Inhibitor Cocktail (Roche, 1187358001) and 2 Unit of MNase (Roche, 10107921001) per million cells. The samples were then incubated with 150 mM NaCl for 10 min at 4 °C. Samples were centrifuged at 16,000 × g for 5 min and the supernatant was collected. 250ug of lysate was precleared with 20 μl of packed Protein A Sepharose CL-4B resin (Cytivia, 17078001) for 2 hours at 4 °C and then incubated overnight with 3ug of anti-NPM1 antibody (Novus Biologicals, NB600-1030) at 4 °C. IgG (millipore, 12-371) was used as a control for IP. Next day, samples were incubated with Protein G Dynabeads (Invitrogen, 10004D) for 2 hours. Beads were then washed 3 times in C-100 buffer (20 mM Hepes pH 7.6, 20% glycerol, 200 mM NaCl, 1.5 mM MgCl2, 0.2 mM EDTA, 0.02% NP40, and 1x cOmplete™ Protease Inhibitor Cocktail). NPM1-interacting proteins on beads were submitted for tryptic digestion off the beads and subsequent mass spectrometric analyses.

G9a-IP was performed on HEK 293T cells. 48 hours post-transfection of the indicated plasmids, cells were washed twice in cold 1X PBS and then incubated with hypotonic buffer (10mM HEPES, 1.5mM MgCl_2_, 10nM KCl, 1x cOmplete™ Protease Inhibitor Cocktail) for 10 minutes on ice. Samples were supplemented with Triton X-100 at a final concentration of 0.5% for 10 minutes on ice. Then, cell nuclei were lysed for 30 minutes at 37°C in MNase digestion buffer (50 mM Tris–HCl pH 7.5, 0.3 M Sucrose, 30 mM KCl, 7.5 mM NaCl, 4 mM MgCl_2_, 1 mM CaCl_2_, 0.125% NP-40) freshly supplemented with 0.25% NaDeoxycholate, 1x cOmplete™ Protease Inhibitor Cocktail and 2 Unit of MNase (Roche, 10107921001) per million cells. The samples were then incubated with 150 mM NaCl for 10 min at 4 °C.

Samples were centrifuged at 16,000 × g for 5 min and the supernatant was collected. 250ug of lysate was precleared with 20 μl of packed Protein A Sepharose CL-4B resin (Cytivia, 17078001) for 2 hours at 4 °C. Samples were then spun down and the cleared supernatant was transferred to a new tube. 20% of the lysate was resuspended in Laemmli loading buffer to use as input while the rest was incubated overnight at 4°C with 20 μl of Anti-HA magnetic beads (Pierce, 88836). Next day, the beads were washed 3 times at 4°C for 10 minutes with C-100 buffer (20 mM HEPES pH 7.6, 20% Glycerol, 200 mM NaCl, 1.5 mM MgCl_2_, 0.2 mM EDTA, 0.02% NP-40), resuspended in Laemmli loading buffer, and incubated at 95°C for 5 minutes. The samples were analysed by SDS-PAGE and Western blotting. The mass spectrometry proteomics data were handled using the local laboratory information management system b-fabric (LIMS) ^45^. The acquired shotgun MS data were processed for identification and quantification using DIA-NN ^46^. Spectra were searched against a Uniprot Homo sapiens reference proteome (reviewed canonical version from 2023-03-30), concatenated to common protein contaminants. The R package prolfqua ^47^ was used to analyze the differential expression and to determine group differences and false discovery rates for all quantifiable proteins. The fold change of average intensities of the 3 replicates of NPM1-IP and IgG control were log_2_ transformed and plotted in prism (Version 9.4.1).

### Quantification and statistical analysis

The number of independent experimental replications is reported in the figure legends. Statistical significance (*P*-values) was calculated using the paired or unpaired two-tailed *t*-test as specified in the corresponding figure legends. For all statistical analyses, a value of *P* < 0.05 was statistically significant.

## Acknowledgements

This work was supported by the ERC grant (ERC-AdG-787074-NucleolusChromatin), Swiss National Science Foundation (31003B-201268), and Swiss Cancer Research Foundation (KFS-5488-02-2022). We thank Catherine Aquino, Tobias Kockmann, and the Functional Genomic Center Zurich for the assistance in sequencing and mass-spectrometry. We thank Antoine Peters for providing the *HP1α/β*-cDKO ESCs.

## Author contributions

S.G. performed and analysed DNA-FISH, IFs, ChIPseq, and RNAseq experiments and established NPM1 cell lines. C.B. performed Nucleolar-DamID and HP1-ChIPseq analyses. M.R. performed DNA-FISH in HP1-KO cells. D.B. cloned all the NPM1 constructs. L.S performed experiments with NSC348884 treatment. All authors contributed to experimental design and data interpretation. R.S conceived and supervised the project.

## Extended Data

**Extended Data Figure 1 NPM1 is required for NAD association with nucleoli**

**a-c.** Representative immuno-FISH images of NADs at chromosome 1 (**a**), 5 (**b**), and 19 (**c**) in ESCs depleted of NPM1 via siRNA.

**d.** Relative expression levels of *NPM1*, Nucleolin (*Ncl*), and Fibrillarin (*Fbl*) in ESCs treated with siRNA-Control and the corresponding siRNAs.

**e, f.** Representative immuno-FISH images for NADs at chromosome 1, 5, and 19 in ESCs depleted of Nucleolin (NCL, **e**) or Fibrillarin (FBL, **f**) via siRNA.

**Extended Data Figure 2 NPM1-KO impairs NAD association with nucleoli**

**a.** Immunofluorescence images showing NPM1 levels in the FKBP12^F36V^-NPM1-ESC line upon treatment with 100 nM dTAG-13 for 72 hours. Nucleolin serves as nucleolar marker.

**b.** Western blot showing NPM1 expression levels upon treatment with dTAG-13 at the indicated concentrations for 72 hours. Using NPM1 and β-actin antibodies. β-actin serves as loading control.

**c.** Representative immuno-FISH images for NADs at chromosome 2 in the FKBP12^F36V^-NPM1-ESC line upon treatment with 100 nM dTAG-13 for 72 hours. Nucleolin serves as nucleolar marker.

**d.** Quantifications of immuno-FISH analyses showing the percentage of FKBP12^F36V^-NPM1-ESCs displaying NADs at the indicated chromosomes that contact nucleoli with or without treatment with 100 nM dTAG-13 for 72 hours. Error bars represent s.d. Statistical significance (*P*-values) from three independent experiments was calculated using the Mann-Whitney test (** < 0.01; *** < 0.001; ns: non-significant).

**e.** Cell cycle analysis of ESCs upon NPM1 depletion. Data are from three independent experiments of FKBP12^F36V^-NPM1-ESC treated with 100 nM dTAG-13 for 72 hours. Error bars represent s.d. Statistical significance (*P*-values) for three independent experiments was calculated using t-test (*< 0.05; ns: non-significant).

**f.** Representative immunofluorescence images of NIH3T3 cells co-transfected with H2B-GFP or H2B-GFP-NoLS expressing plasmids shRNA-Control or shRNA-*NPM1* sequences. Bar represents 5 μm.

**Extended Data Figure 3 HP1 is not required for NAD association with nucleoli**

**a.** Representative immunofluorescence images of *HP1 α/β*-cDKO ESCs treated without and with 1 μM 4-OHT for 48 hours.

**b.** Representative immuno-FISH images for NADs at chromosome 2 and 19 in *HP1 α/β*-cDKO ESCs without and with 1 μM 4-OHT for 48 hours.

**c.** Quantifications of immuno-FISH analyses showing the percentage of cells displaying NADs contacting nucleoli in *HP1 α/β*-cDKO ESCs upon treatment with 4-OHT. Data are from two independent experiments. Mean values are shown.

**d.** Measurement of HP1α, β, and γ genome occupancy at the active A compartment, NAD-only, and NAD/LAD regions in ESCs. Data are from ^26^. Values are shown as average RPKM of a 10kb bin size region. Error bars represent s.d.. Statistical significance (*P*-values) was calculated using the unpaired two-tailed t test (****<0.0001).

**e.** Representative immunofluorescence images showing H3K9me3 distribution in ESCs treated with siRNA-Control or siRNA-*NPM1*. UBF serves as nucleolar marker.

**f.** DAPI-stained images of ESCs treated with siRNA-Control or siRNA-*NPM1* shown in Figure 3A.

**g.** Representative immuno-FISH images for major satellites in FKBP12^F36V^-NPM1-ESCs upon treatment with 100 nM dTAG-13 for 72 hours. Nucleolin serves as nucleolar marker

**Extended Data Figure 4 G9a is not required for NAD association with nucleoli**

**a.** Biological functions and cellular components of NPM1-interacting proteins detected by String analyses.

**b.** Relative expression levels of G9a upon G9a-KD measured by qRT-PCR. Data are from three independent experiments. Error bars represent s.d.. Statistical significance (*P*-values) was calculated using the unpaired two-tailed t test (**<0.01).

**c.** Western blot showing global H3K9me2 levels in ESCs upon G9a depletion. Tubulin serves as loading control.

**d.** Top 10 gene ontology (GO) terms as determined using DAVID for genes upregulated and downregulated upon G9a-KD.

**e.** Representative immuno-FISH images of NADs at chromosome 2 in ESCs depleted of G9a-KD.

**f.** Quantifications of immuno-FISH analyses showing the percentage of cells displaying NADs contacting nucleoli in ESCs depleted of G9a. Error bars represent s.d. Statistical significance (*P*-values) from three independent experiments was calculated using the Mann-Whitney test (ns: non-significant).

**g.** Relative resistance to DpnII digestion in ESCs expressing H2B-Dam-NoLS-DD transfected with siRNA-Control or with siRNA-G9a. Expression of H2B-Dam-NoLS-DD was induced by treatment with doxycycline and Shield for 15 hours. Data are from two independent experiments. The mean values are shown.

**Extended Data Figure 5 NPM1 oligomerization is not required for NAD association with nucleoli**

**a.** Western blots showing NPM1 signals upon treatment for 24 hours with NSC348884 at the indicated concentrations. Upper panel, native protein gel. Lower panel, denaturating protein gel.

**b.** Representative immunofluorescence images showing NPM1 in ESCs treated with for 24 hours with 0.5 μM NSC348884.

**c.** Representative immuno-FISH images of NADs at chromosome 1 in ESCs treated for 24 hours with 0.5 μM NSC348884.

