## Supplementary Figures for "NPM1 mediates genome-nucleolus interactions and the establishment of their repressive chromatin states"

### Extended Data Figure 1

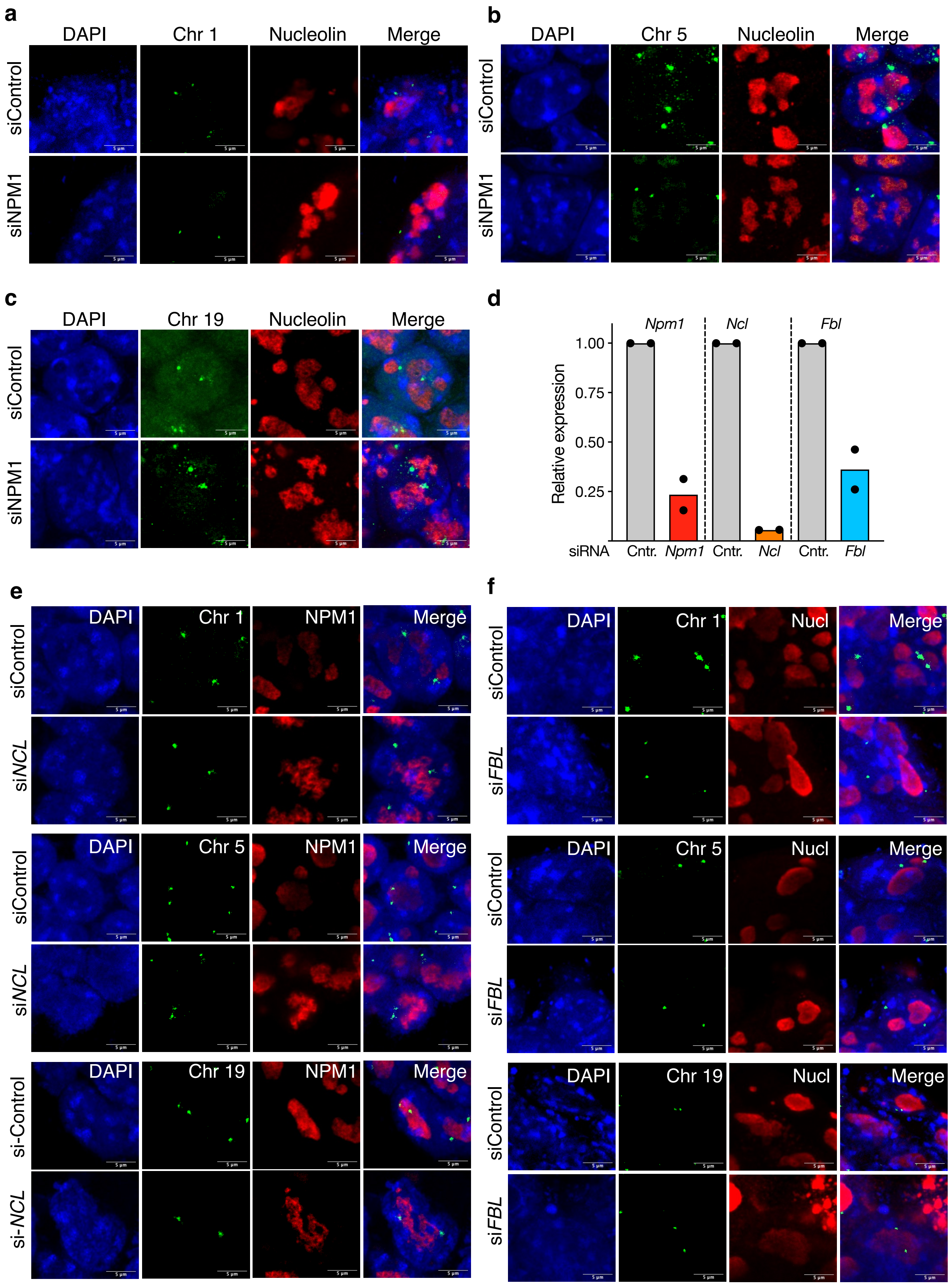

### Extended Data Figure 2

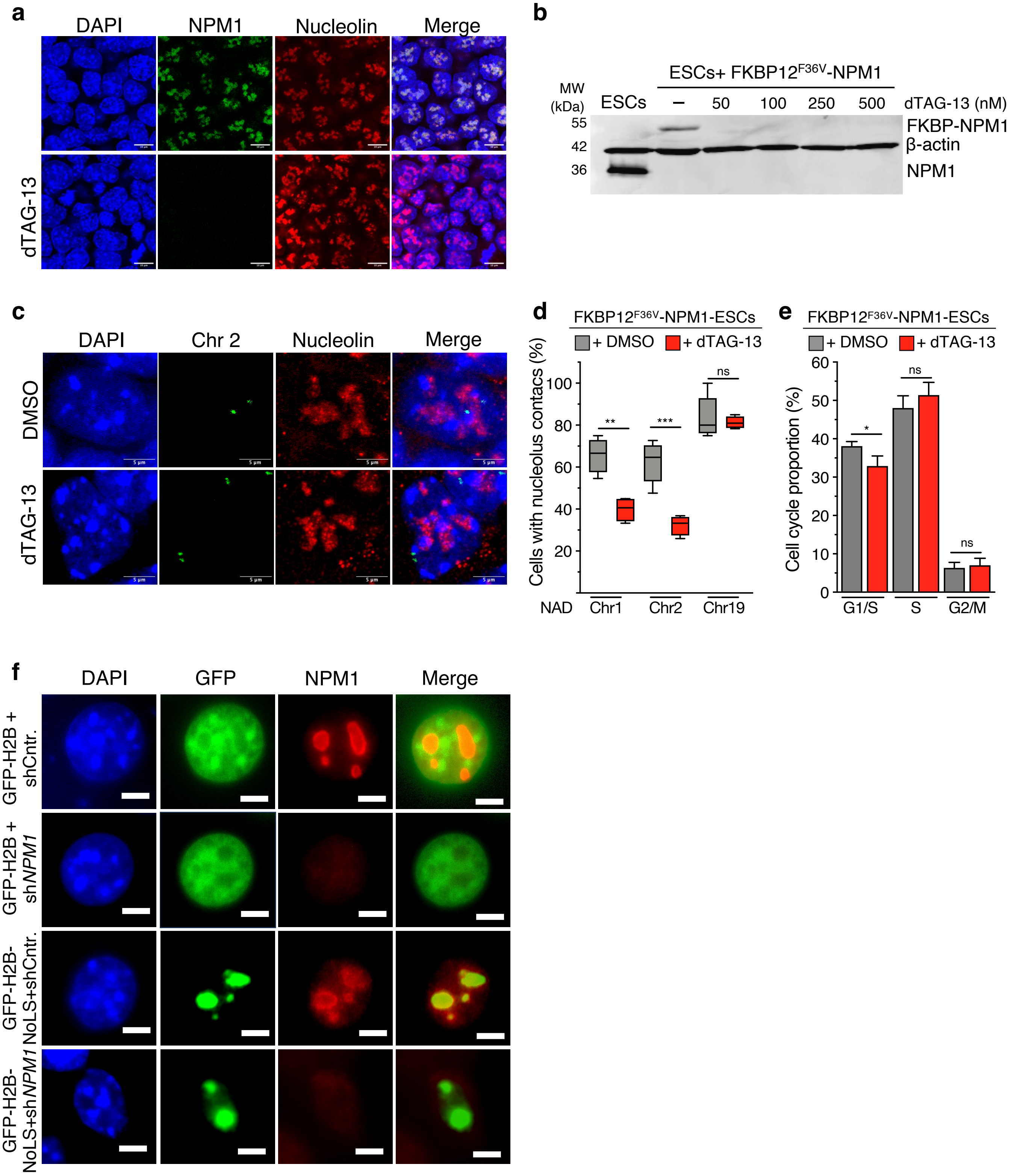

### Extended Data Figure 3

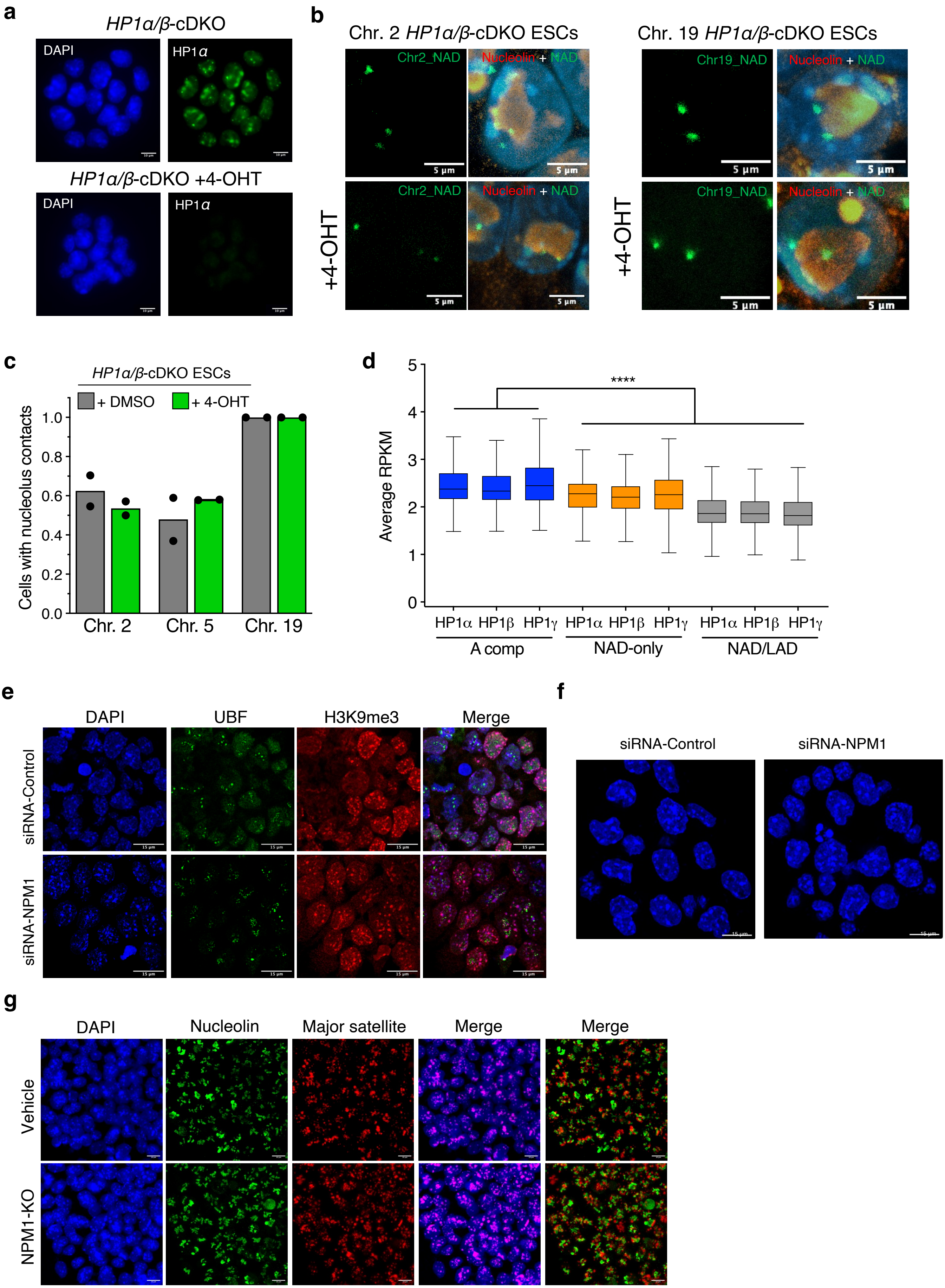

### Extended Data Figure 4

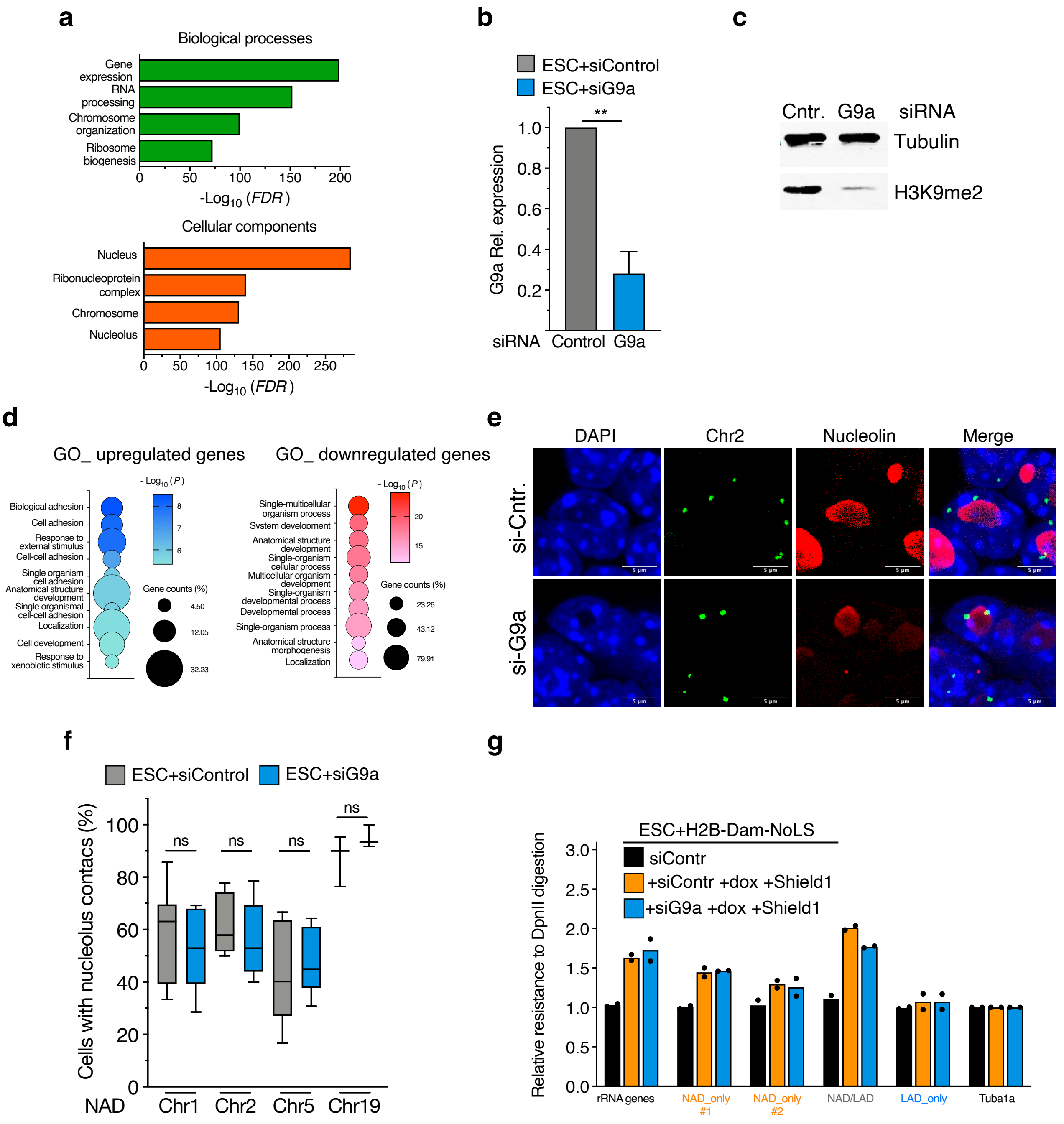

### Extended Data Figure 5

**a**

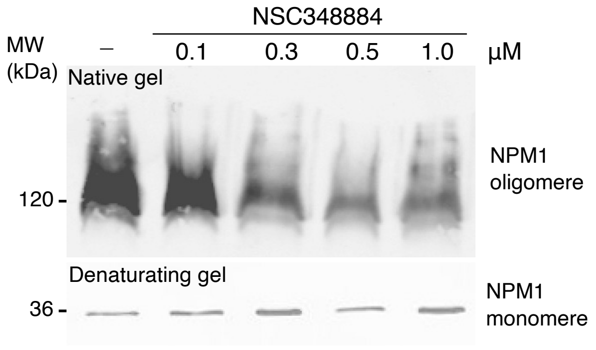


**b**

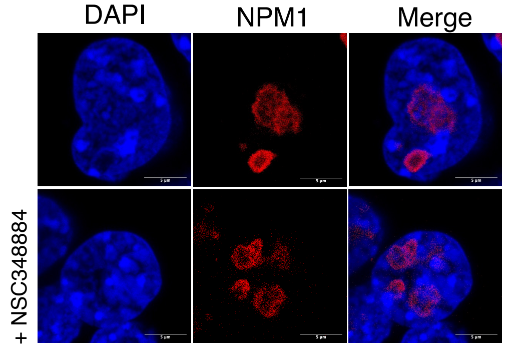


**c**

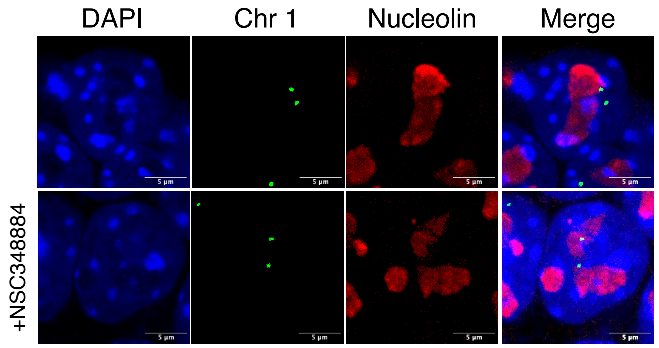
